# Genetic regulatory mechanisms of smooth muscle cells map to coronary artery disease risk loci

**DOI:** 10.1101/309559

**Authors:** Boxiang Liu, Milos Pjanic, Ting Wang, Trieu Nguyen, Michael Gloudemans, Abhiram Rao, Victor G. Castano, Sylvia Nurnberg, Daniel J. Rader, Susannah Elwyn, Erik Ingelsson, Stephen B. Montgomery, Clint L. Miller, Thomas Quertermous

## Abstract

Coronary artery disease (CAD) is the leading cause of death globally. Genome-wide association studies (GWAS) have identified more than 95 independent loci that influence CAD risk, most of which reside in non-coding regions of the genome. To interpret these loci, we generated transcriptome and whole-genome datasets using human coronary artery smooth muscle cells (HCASMC) from 52 unrelated donors, as well as epigenomic datasets using ATAC-seq on a subset of 8 donors. Through systematic comparison with publicly available datasets from GTEx and ENCODE projects, we identified transcriptomic, epigenetic, and genetic regulatory mechanisms specific to HCASMC. We assessed the relevance of HCASMC to CAD risk using transcriptomic and epigenomic level analyses. By jointly modeling eQTL and GWAS datasets, we identified five genes (*SIPA1*, *TCF21*, *SMAD3*, *FES*, and *PDGFRA*) that modulate CAD risk through HCASMC, all of which have relevant functional roles in vascular remodeling. Comparison with GTEx data suggests that *SIPA1* and *PDGFRA* influence CAD risk predominantly through HCASMC, while other annotated genes may have multiple cell and tissue targets. Together, these results provide new tissue-specific and mechanistic insights into the regulation of a critical vascular cell type associated with CAD in human populations.

## Introduction

Atherosclerotic coronary artery disease (CAD) is the leading cause of death in both developed and developing countries worldwide, and current estimates predict that more than 1 million individuals will suffer from new and recurrent CAD this year in the U.S. alone^1^. Like most polygenic diseases, both genetic and environmental factors influence an individual’s lifetime risk for CAD^2^. Early Swedish twin studies and more recent genome-wide association studies (GWAS) have estimated that about 50% of CAD risk is explained by genetic factors^3,4^. To date, GWAS have reported more than 95 replicated independent loci and numerous additional loci that are associated at an FDR<0.05^5–8^. A majority of these loci reside in non-coding genomic regions and are expected to function through regulatory mechanisms. Also, approximately 75% of CAD loci are not associated with classical risk factors, suggesting that at least part of them function through mechanisms intrinsic to the vessel wall.

Smooth muscle cells (SMC) constitute the majority of cells in the coronary artery wall. In response to vascular injury (e.g. lipid accumulation, inflammation), SMCs undergo phenotypic switching, and ultimately contribute to both atherosclerotic plaque formation and stabilization^9–12^. Recent lineage tracing studies in mice have revealed that although 80% of plaque-derived cells lack traditional SMC markers, roughly half are of SMC origin^13,14^. Thus, genetic studies of human coronary artery smooth muscle cells (HCASMC) have the potential to shed new light on their diverse functions in the vessel wall relevant to human atherosclerosis. In a few cases, the underlying mechanisms have been identified for CAD loci in vascular SMC models^10,15–18^. Large-scale expression quantitative trait loci (eQTL) mapping efforts such as the Genotype Tissue Expression (GTEx) project have helped refine these mechanisms for multiple traits across human tissues^19^. However, due to the lack of HCASMC in both GTEx and other studies, the overall contribution of this cell type towards heritable CAD risk remains unknown.

Herein, we performed whole-genome sequencing and transcriptomic profiling of 52 HCASMC donors to quantify the effects of *cis*-acting variation on gene expression and splicing associated with CAD. We evaluated the tissue specificity and disease relevance of our findings in HCASMC by comparing to publicly available GTEx and ENCODE datasets. We observed significant colocalization of eQTL and GWAS signals for five genes (*FES*, *SMAD3*, *TCF21*, *PDGFRA* and *SIPA1*), which all have the capacity to perform relevant functions in vascular remodeling. Further, comparative analyses with GTEx datasets reveals that *SIPA1* and *PDGFRA* act primarily in HCASMC. Together, these findings demonstrate the power of leveraging genetics of gene regulation for a critical cell type to uncover new risk-associated mechanisms for CAD.

## Material and Methods

### Sample acquisition and cell culture

A total of 62 primary human coronary artery smooth muscle cell (HCASMC) lines collected from donor hearts were purchased, and 52 lines remained after stringent filtering (see **Supplementary Note**). These 52 lines were from PromoCell (catalog # C-12511, n = 19), Cell Applications (catalog # 350-05a, n = 25), Lonza (catalog # CC-2583, n = 3), Lifeline Cell Technology (catalog # FC-0031, n = 3), and ATCC (catalog # PCS-100-021, n = 2). All lines were stained with smooth muscle alpha actin to check for smooth muscle content and all lines tested negative for mycoplasma (Table S1). All cell lines were cultured in smooth muscle growth medium (Lonza catalog # CC-3182) supplemented with hEGF, insulin, hFGF-b, and 5% FBS, according to Lonza instructions. All HCASMC lines were expanded to passage 5-6 prior to extraction.

### Library preparation and sequencing

<u>Whole Genome Sequencing:</u> Genomic DNA was isolated using Qiagen DNeasy Blood & Tissue Kit (catalog # 69506) and quantified using NanoDrop 1000 Spectrophotometer (Thermo Fisher). Macrogen performed library preparation using Illumina’s TruSeq DNA PCR-Free Library Preparation Kit and 150 bp paired-end sequencing on Illumina HiSeq X Ten System. <u>RNA Sequencing:</u> RNA was extracted using Qiagen miRNeasy Mini Prep Kit (catalog # 74106). Quality of RNA was assessed on the Agilent 2100 Bioanalyzer. Samples with RIN greater than or equal to 8 were sent to the Next-Generation Sequencing Core at the Perelman School of Medicine at the University of Pennsylvania. Libraries were made using Illumina TruSeq Stranded Total RNA Library Prep Kit (catalog # 20020597) and sequenced using 125bp paired-end on HiSeq 2500 Platform. <u>ATAC Sequencing:</u> We used ATAC-seq to assess chromatin accessibility with slight modifications to the published protocol^57^. Approximately 5m10^4^ fresh cells were collected at 500 g, washed in PBS, and nuclei extracted with cold lysis buffer. Pellets were subjected to transposition containing Tn5 transposases (Illumina) at 37 °C for 30 min, followed by purification using the DNA Clean-up and Concentration kit (Zymo). Libraries were PCR amplified using Nextera barcodes, with the total number of cycles empirically determined using SYBR qPCR. Amplified libraries were purified and quantified using bioanalyzer, nanodrop and qPCR (KAPA) analysis. Libraries were multiplexed and 2×75 bp sequencing was performed using an Illumina NextSeq 500.

### Alignment and quantification of genomic, transcriptomic and epigenomic features

Whole-genome sequencing data were processed with the GATK best practices pipeline with hg19 as the reference genome^20,58^, and VCF records were phased with Beagle v4.1^59^. Variants with imputation allelic r^2^ less than 0.8 and Hardy-Weinberg Equilibrium p-value less than 1×10^−6^ were filtered out (see **Supplementary Note**). Demultiplexed FASTQ files were mapped with STAR version 2.4.0i in 2-pass mode^60^ over the hg19 reference genome. Prior to expression quantification, we filtered our reads prone to mapping bias using WASP^61^. Total read counts and RPKM were calculated with RNA-SeQC v1.1.8^62^ using default parameters with additional flags “-n 1000 -noDoC -strictMode” over GENCODE v19 reference. Allele-specific read counts were generated with the createASVCF module in RASQUAL^27^. We quantified intron excision levels using LeafCutter intron-spanning reads^63^. In brief, we converted bam files to splice junction files using the bam2junc.sh script, and defined intron clusters using leafcutter_cluster.py with default parameters, which requires at least 30 reads supporting each intron and allows intron to have a maximum size of 100kb. We used the ENCODE ATAC-seq pipeline to perform alignment and peak calling (https://github.com/kundajelab/atac_dnase_pipelines)64. FASTQ files were trimmed with Cutadapt v1.9^65^ and aligned with Bowtie2 v2.2.6^66^. MACS2 v2.0.8^67^ was used to call peaks with default parameters. Irreproducible Discovery Rate (IDR)^68^ analyses were performed based on pseudo-replicates (subsample of reads) with a cutoff of 0.1 to output an IDR call set, which was used for downstream analysis. We used WASP^61^ to filter out reads that are prone to mapping bias.

### Mapping of cis-acting quantitative trait loci (QTL)

Prior to QTL mapping, we inferred ancestry principal components (PCs) using the R package SNPRelate^69^ on a pruned SNP set (Fig. S4). We filtered out SNPs based on Hardy-Weinberg equilibrium (HWE < 1×10^−6^), LD (r^2^ < 0.2) and minor allele frequency (MAF < 0.05)^69^. To correct for hidden confounders, we extracted 15 covariates using PEER^70^ on quantile normalized and rank-based inverse normal transformed RPKM values. The number of hidden confounders to removed was determined by empirically maximize the power to discover eQTLs on chromosome 20 (for computational speed and to avoid overfitting). We tested combinations of 3 to 5 genotype principal components with 1 to 15 PEER factors. We found that the combination of 4 genotype PCs with 8 PEER factors provides the most power to detect eQTLs. We then used sex, the top four genotype principal components, and the top eight PEER factors in both FastQTL and RASQUAL to map cis-eQTL with a 2Mb window centered at transcription start sites. Mathematically, the model is the following:

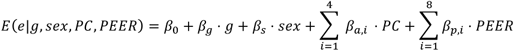

Where *e* stands for gene expression, and *g* stands for the genotype of the test SNP. We used LeafCutter^31^ to quantify intron excision levels, and FastQTL^26^ to map cis-sQTLs within a 200 kbp window around splice donor sites, controlling for sex, genotype PCs, and splicing PCs. Using a similar approach, we found that 3 genotype PCs and 6 splicing PCs maximized the power to map sQTLs. To control for multiple hypothesis testing, we calculated per-gene eQTL p-values using FastQTL with permutation, and controlled transcriptome-wide false discovery rate with the q-value package^71^. For RASQUAL, it was not computationally feasible to perform gene-level permutation testing. Instead, we used TreeQTL to simultaneously control for SNP-level FDR and gene-level FDR^72^. Note that TreeQTL is more conservative than permutation {Consortium:jn}.

### Quantifying tissue- and cell-type specific contribution to coronary artery disease (CAD) risk

We used stratified LD score regression^33^ to estimate the enrichment of heritability for SNPs around tissue- and cell-type specific genes as described previously^32^. We defined tissue-specific genes by first selecting independent tissues and removing tissues primarily composed of smooth muscle to avoid correlation with HCASMC (see **Supplementary note**). After filtering, 16 tissues remained: HCASMC, Adipose - Subcutaneous, Adrenal Gland, Artery - Coronary, Brain - Caudate (basal ganglia), Cells - EBV-transformed lymphocytes, Cells - Transformed fibroblasts, Liver, Lung, Minor Salivary Gland, Muscle - Skeletal, Pancreas, Pituitary, Skin - Not Sun Exposed (Suprapubic), Testis, Whole Blood. We defined tissue-specific genes using gene expression score. For each gene, we determined the mean and standard deviation of median RPKM across tissues, from which the z-score is derived.

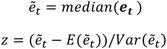

Where ***e_t_*** is the RPKM across all individuals in tissue *t*. We ranked each gene based on the z-scores (a higher z-score indicates more tissue specificity), and defined tissue-specific genes as the top 1000, 2000, and 4000 genes. A given SNP was assigned to a gene if it fell into the union of exon +/− 1kbp of that gene. We estimated the heritability enrichment using stratified LD score regression on a joint SNP annotation across all 16 tissues against the CARDIoGRAMplusC4D GWAS meta-analysis^73^. To determine whether CAD risk variants are enriched in the open chromatin regions tissue- and cell-type specific fashion, we used a modified version of GREGOR^34^ to estimate the likelihood of observing given number of GWAS variants falling into open chromatin regions of each tissue and cell type (see **Supplementary Note**). We first defined a GWAS locus as all variants in LD (r^2^>0.7) with the lead variant. Given a set of GWAS loci, we selected 500 background variants matched by 1) number of variants in LD, 2) distance to the nearest gene, and 3) minor allele frequency, and 4) gene density in a 1Mb window. We calculated p-values and odds ratios between GWAS variants and background variants across HCASMC and all ENCODE tissues and primary cell lines.

### Colocalization between molecular QTL and CAD genome-wide association study (GWAS)

We used summary-data-based Mendelian Randomization (SMR)^37^ to determine GWAS loci that can be explained by *cis-*acting QTLs. We performed colocalization tests for 3,379 genes with cis-eQTL p-value < 5×10^−5^ for the top variant and 2,439 splicing events with cis-sQTL p-value < 5×10^−5^ for the top variant in HCASMC against the latest CARDIoGRAMplusC4D and UK Biobank GWAS meta-analysis^6^. We identified genome-wide significant eQTL and sQTL colocalizations based on adjusted SMR p-values (Benjamini-Hochberg FDR < 0.05). The equivalent p-value was 2.96×10^−5^ and 2.05×10^−5^ for eQTL and sQTL, respectively. SMR uses a reference population to determine linkage between variants; we used genetic data from individuals of European ancestry from 1000 Genomes as the reference population in our analyses. We also used a modified version of eCAVIAR^35^ to identify colocalized signals (see **Supplementary Note**). We calculated colocalization posterior probability (CLPP) using all SNPs within 500kb of the lead eQTL SNP against CAD summary statistics from CARDIoGRAMplusC4D and UK Biobank GWAS meta-analysis^6^. For computational feasibility, the GWAS and eQTL loci were assumed to have exactly one causal SNP. We defined colocalization events using CLPP > 0.05. Note that this is more conservative than the default eCAVIAR cutoff (CLPP > 0.01). We determined the direction of effect, namely whether gene upregulation increases risk, using the correlation of effect sizes in the GWAS and the eQTL studies. We selected SNPs with p-value < 1×10^−3^ in both the GWAS and eQTL datasets (since other SNPs carry mostly noise), and fitted a regression using the GWAS and eQTL effect sizes as the predictor and the response, respectively. We defined the direction of effect as the sign of the regression slope.

## Results

### HCASMC-specific genomic architecture

We obtained and cultured 62 primary HCASMC lines, and 52 lines remained for analysis after stringent quality control (**Supplementary Note** and Table S1). We performed whole-genome sequencing to an average depth of 30X, and jointly called genotypes using the GATK best practices pipeline^20^, producing a total of ~15.2 million variants after quality control (see **Methods**). For RNA, we performed 125bp paired-end sequencing to a median depth of 51.3 million reads, with over 2.7 billion reads in total. After quantification and quality control, 19,607 genes were expressed in sufficient depth for downstream analysis (Table 1). To confirm that HCASMC derived from tissue culture reflect *in vivo* physiology, we first projected their transcriptomes onto the 53 tissues profiled in GTEx^19^ (Fig. 1A). Using multi-dimensional scaling (MDS) to visualize the similarity of HCASMC to GTEx tissues, we observed that HCASMC forms a distinct cluster and closely neighbors fibroblasts, skeletal muscle, arteries, heart and various smooth muscle-enriched tissues (vagina, colon, stomach, uterus and esophagus). These results were expected given that HCASMC are predicted to be similar to skeletal muscle, smooth muscle-enriched tissues as well as tissues representing the same anatomical compartment (e.g. heart and artery)^21^. In addition, HCASMC resemble fibroblast as both can be differentiated from mesenchymal cells from the dorsal mesocardium^22^. We also computed the epigenetic similarity between HCASMC and ENCODE cell types^23^. Consistent with the transcriptomic findings, the closest neighbors to HCASMC using epigenomic data were fibroblasts, heart, lung and skeletal muscle (Fig. 1B).

**Table 1.** Molecular quantitative trait loci discoveries. We report the number of tests performed and the number of significant loci at FDR < 0.05, 0.01, and 0.001 for eQTL and sQTL stratified by molecular trait type. We used permutation and the Benjamini-Hochberg adjustment for sQTL discovery, and a multi-level FDR correction procedure (TreeQTL^72^) for eQTL discovery, where permutation was not computationally feasible (see **Methods**).

| Molecular Phenotype | Trait type | # of traits tested | # of traits with at least one QTL |  |  |
| --- | --- | --- | --- | --- | --- |
|  |  |  | FDR = 0.05 | FDR= 0.01 | FDR = 0.001 |
| Gene expression | Protein coding | 13624 | 1048 (7.69%) | 841 (6.17%) | 636 (4.67%) |
|  | lincRNA | 1266 | 51 (4.03%) | 41 (3.24%) | 33 (2.61%) |
|  | Pseudogene | 2616 | 50 (1.91%) | 34 (1.3%) | 25 (0.96%) |
|  | Other | 2101 | 71 (3.38%) | 56 (2.67%) | 44 (2.09%) |
|  | Total | 19607 | 1220 (6.22%) | 972 (4.96%) | 738 (3.76%) |
| Splicing | Protein coding | 24461 | 519 (2.12%) | 349 (1.43%) | 245 (1%) |
|  | lincRNA | 300 | 11 (3.67%) | 7 (2.33%) | 5 (1.67%) |
|  | Pseudogene | 376 | 22 (5.85%) | 15 (3.99%) | 12 (3.19%) |
|  | Other | 541 | 29 (5.36%) | 19 (3.51%) | 17 (3.14%) |
|  | Total | 25678 | 581 (2.96%) | 390 (1.99%) | 279 (1.42%) |

**Fig. 1.**
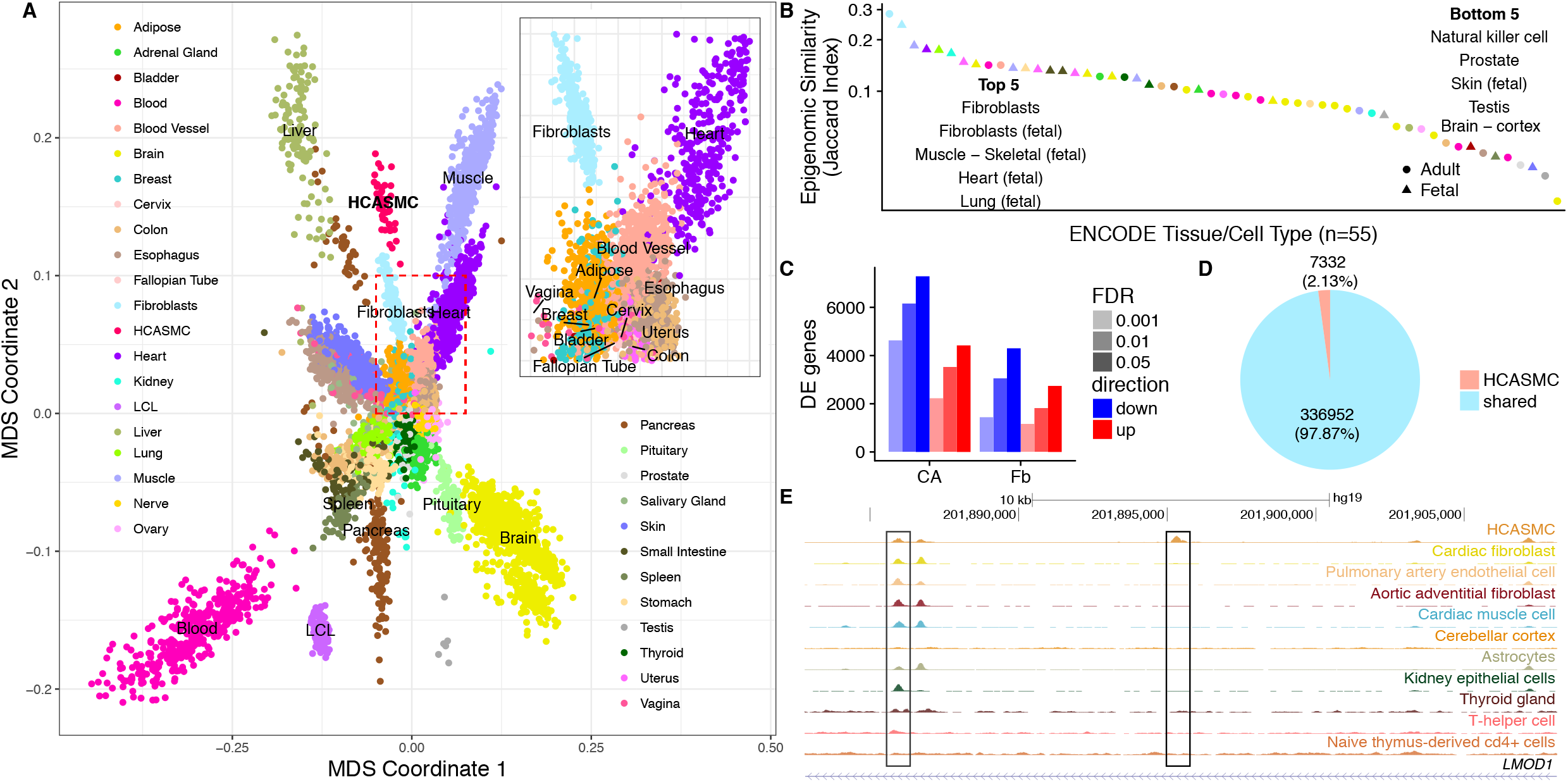
The relationship between HCASMC and GTEx and ENCODE cell and tissue types. (**A**) The multidimensional scaling plot of gene expression shows that HCASMC form a distinct cluster, which neighbors fibroblast, skeletal muscle, heart, blood vessel and various types of smooth muscle tissues such as esophagus and vagina (inset). (**B**) Jaccard similarity index between HCASMC and ENCODE cell and tissue types reveals that fibroblast, skeletal muscle, heart and lung are most closely related to HCASMC. (**C**) Thousands of genes are differentially expressed between HCASMC and its close neighbors, fibroblast, as well as the tissue of origin, coronary artery. (**D**) A total of 344284 open chromatin peaks are found in HCASMC, of which 7332 (2.1%) are HCASMC-specific. (**E**) An example of a HCASMC-specific peak located within the intron of *LMOD1*, which is an HCASMC-specific gene.

Next, we determined the pathways that may be selectively upregulated in HCASMC compared to closely related tissues. We performed differential expression analysis of HCASMC against fibroblasts and coronary artery in GTEx after correcting for batch effects and other hidden confounders (see **Methods**). Overall, 2,610 and 6,864 genes were found to be differentially expressed, respectively (FDR < 1×10^−3^, Fig. 1C and Fig. S1), affecting pathways involved in cellular proliferation, epithelial-mesenchymal transition (EMT) and extracellular matrix (ECM) secretion (Table S2). Next, we sought to identify HCASMC-specific epigenomic signatures by comparing HCASMC open chromatin profiles, as determined with ATAC-seq, against DNaseI hypersensitivity (DHS) sites across all ENCODE primary cell types and tissues (Table S3). We processed HCASMC ATAC-seq data with the ENCODE pipeline and standardized peaks as 75 bp around the peak summit for all tissues and cell lines to mitigate batch effect (see **Methods**). A total of 7332 peaks (2.1%) were not previously identified in ENCODE and represent HCASMC-specific sites (Fig. 1D). For example, an intronic peak within the *LMOD1* gene was found to be unique to HCASMC (Fig. 1E). This gene is expressed only in vascular and visceral smooth muscle cells where it is involved in actin polymerization, and has been mapped as a candidate causal CAD gene^11^. We then sought to identify transcription factor binding sites overrepresented in HCASMC-specific peaks. Motif enrichment analyses indicated that HCASMC-specific open chromatin sites are enriched with binding sites for members of the forkhead box (FOX) transcription factor family (see **Methods**). We performed motif enrichment analysis using 50-, 200-, and 1000-bp regions flanking HCASMC-specific peaks, and found that the enrichment was robust to selection of window size, indicating the result is not simply due to selection bias (Fig. S2). The FOX transcription factors are known to regulate tissue- and cell-type specific gene transcription^24^, and a subgroup of this family includes those with the ability to serve as pioneer factors^25^. To validate that FOX motif enrichment is specific to HCASMC, we performed similar analyses for brain-, heart-, and fibroblast-specific open chromatin regions and observed a depletion of FOX motifs (Fig. S3). Together these results suggest that HCASMC-specific transcriptomic and epigenomic profiles provide new regulatory mechanisms previously lacking in large publicly available datasets.

### Expression and splicing quantitative trait locus discovery

In order to investigate the genetic regulatory mechanisms of gene expression in HCASMC, we conducted genome-wide mapping of eQTLs using both FastQTL^26^ and RASQUAL^27^ on the 52 donor samples from diverse ethnic backgrounds (Table S1 and Fig S4). RASQUAL has been previously shown to increase the cis-eQTL discovery power in small sample sizes by leveraging allele-specific information^27^. Indeed, using a threshold of FDR < 0.05, RASQUAL increased the number of eQTLs discovered approximately seven-fold as compared to FastQTL (RASQUAL:1220 vs. FastQTL:167, Table 1). We next evaluated whether these eQTLs were enriched in regions of open chromatin using data from a subset of individuals with ATAC-seq profiles. We observed that eQTLs within HCASMC open chromatin regions had more significant p-values compared to all eQTLs (Fig. S5, two-sided rank-sum test p-value < 9.2×10^−5^). This is consistent with putative effects of cis-acting variation, potentially functioning through altered TF binding around these accessible regions. Next, using a Bayesian meta-analytic approach^28^, we sought to identify HCASMC-specific eQTLs using GTEx tissues as a reference. Under the most stringent criteria (eQTL posterior probability > 0.9 for HCASMC and < 0.1 for all GTEx tissues, see **Methods**), we identified four HCASMC-specific eQTLs (Fig. S6). For example, rs1048709 is the top eQTL-SNP and confers HCASMC-specific regulatory effects on Complement Factor B (Fig. S6B), a gene that has been previously implicated in atherosclerosis and other inflammatory diseases^29^. In addition to regulatory effects on gene expression, previous studies have identified splicing as a major source of regulatory impact of genetic variation on complex diseases^30^. Therefore, we mapped splicing QTLs (sQTLs) using LeafCutter^31^ and identified 581 sQTLs associated at FDR < 0.05 (Table 1). As a quality control, we estimated the enrichment of sQTLs and eQTLs against a matched set of background variants. As expected, eQTLs were enriched around the 5’ UTR (Fig. S7A), whereas sQTLs were enriched in splicing regions, particularly splice donor and acceptor sites (Fig. S7B).

### Overall CAD genetic risk mediated by HCASMC

We next examined the heritable contribution of HCASMC towards the risk of CAD. Previous reports have suggested that disease-associated SNPs are often enriched in genes expressed in the relevant tissue types^32^. Thus, we estimated the contribution to CAD risk from SNPs in or near genes showing patterns of tissue-specific expression and identified the top 2000 tissue-specific genes for HCASMC and GTEx tissues (see **Methods**). We then applied stratified LD score regression^33^ to estimate CAD heritability explained by SNPs within 1kb of tissue-specific genes. We found that HCASMC, along with coronary artery and adipose tissues, contribute substantially towards CAD heritability (Fig. 2A). These enrichment results were robust to the tissue-specificity cutoff (top 1000, 2000, or 4000 genes), suggesting that they were not simply due to selection bias (Fig. S8). Complementary epigenomic evidence previously demonstrated that risk variants for complex diseases are often enriched in open chromatin regions in relevant tissue types^23,33,34^. Thus, we estimated the degree of overlap between CAD variants and open chromatin in HCASMC and ENCODE cell types using a modified version of GREGOR^34^ (see Methods). We observed that open chromatin regions in HCASMC, as well as vascular endothelial cells, monocytes, uterus (smooth muscle) and B-cells, are enriched for CAD risk variants (Fig. 2B). These findings support the role of HCASMC as an appropriate cellular model to map the genetic basis of CAD, which may be supplemented by the contribution of other vessel wall cell types.

**Fig. 2.**
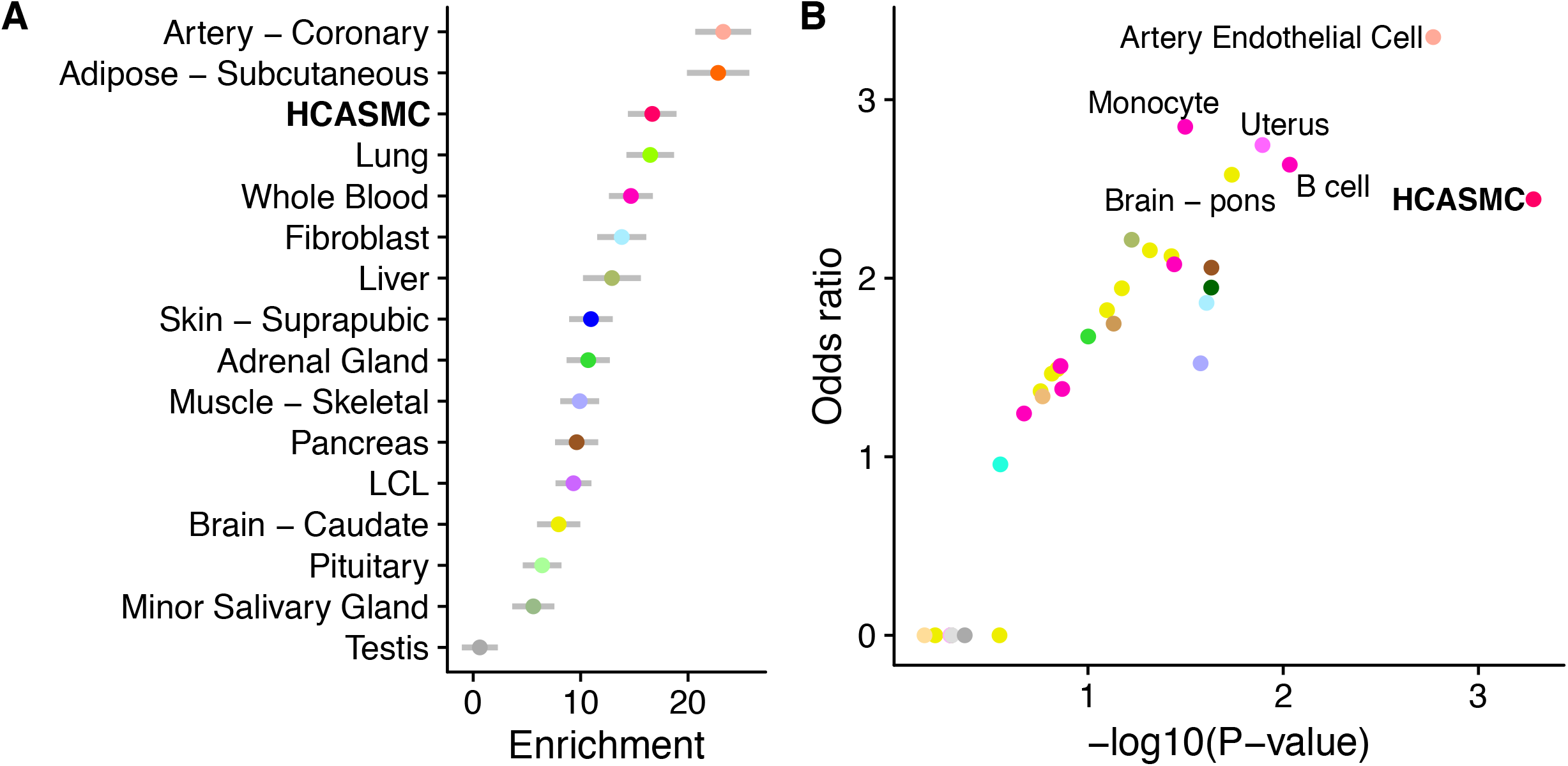
Tissue- and cell-type specific contribution to CAD risk. (**A**) Tissue-specific enrichment of CAD heritability. We used stratified LD score regression to estimate the CAD risk explained by SNPs close to tissue- specific genes, defined as the 2000 genes with highest expression z-scores (see Methods). Genes whose expression is specific to coronary artery, adipose, and HCASMC harbors SNPs with large effects on CAD. (**B**) Overlap between CAD risk variants and tissue- and cell-type specific open chromatin regions. We used a modified version of GREGOR (see **Methods**) to estimate the probability and odds ratio (compared with matched background SNPs) of overlap between CAD risk variants and open chromatin regions in HCASMC and across ENCODE tissues. HCASMC, arterial endothelial cells, monocytes, B cell, uterus (composed primarily of smooth muscle), and pons (possibly through regulation of blood pressure) showed the highest degrees of overlap.

### Fine-mapping CAD risk variants

Whole-genome sequencing of our HCASMC population sample provides the opportunity to fine-map CAD risk loci. Several studies have used colocalization between GWAS and eQTL signals as a fine-mapping approach to identify candidate causal regulatory variants^35–38^, and in several cases pinpointing single causal variants^39,40^. Given the global overlap between CAD risk variants and genetic regulation in HCASMC, we sought to find evidence for colocalization between GWAS and eQTL signals. We thus compiled publicly available genome-wide summary statistics from the latest meta-analysis^6^. We then applied two methods with different statistical assumptions, eQTL and GWAS CAusal Variants Identification in Associated Regions (eCAVIAR)^35^ and Summary-data-based Mendelian Randomization (SMR)^37^ to identify colocalizing variants and genes across all CAD loci, and focused on the union of results from the two independent methods. We used FDR < 0.05 and colocalization posterior probability (CLPP) > 0.05 as cutoffs for SMR and eCAVIAR, respectively (Note that CLPP > 0.05 is more conservative than the CLPP > 0.01 recommended in the publication of the eCAVIAR method). From this approach, we identified five high-confidence genes, namely *FES*, *SMAD3*, *TCF21*, *PDGFRA* and *SIPA1* (Fig. 3). Although the top genes found by two methods differed, we observed that the SMR p-values and eCAVIAR CLPPs positively correlate (Fig. S9), and that two of the three genes found only by eCAVIAR achieved nominal significance in SMR (Table S4). We then investigated whether these colocalizations were unique to HCASMC by conducting colocalization tests across all GTEx tissues. For *SIPA1* and *PDGFRA*, colocalization appears to be HCASMC-specific (Fig. 3G; Fig. S10A and S10D). For *SMAD3*, both HCASMC and thyroid have strong colocalization signals (Fig. S10B). *TCF21* and *FES* colocalization were found to be shared across multiple tissues (Fig. S10C and Fig. S11D). Next, we conducted colocalization analysis between sQTL and GWAS summary statistics with both eCAVIAR and SMR. We identified colocalization with four genes (Table S4, and Fig. S12). The most significant colocalization event is at the *SMG9* locus. Interestingly, the top sQTL variant, rs4760, is a coding variant located in the exon of the *PLAUR* (plasminogen activator urokinase receptor) gene and is also a GWAS variant for circulating cytokines and multiple immune cell traits^41,42^. By correlating eQTL and GWAS effect sizes, we observed that increased *TCF21* and *FES* expression levels are associated with reduced CAD risk, while increased *PDGFRA*, *SIPA1*, and *SMAD3* expression levels are associated with increased CAD risk (Fig. 4A-E). These results provide genetic evidence that pathways promoting SMC phenotypic transition during atherosclerosis can be both protective and detrimental depending on the genes implicated (Fig. 4F).

**Fig. 3.**
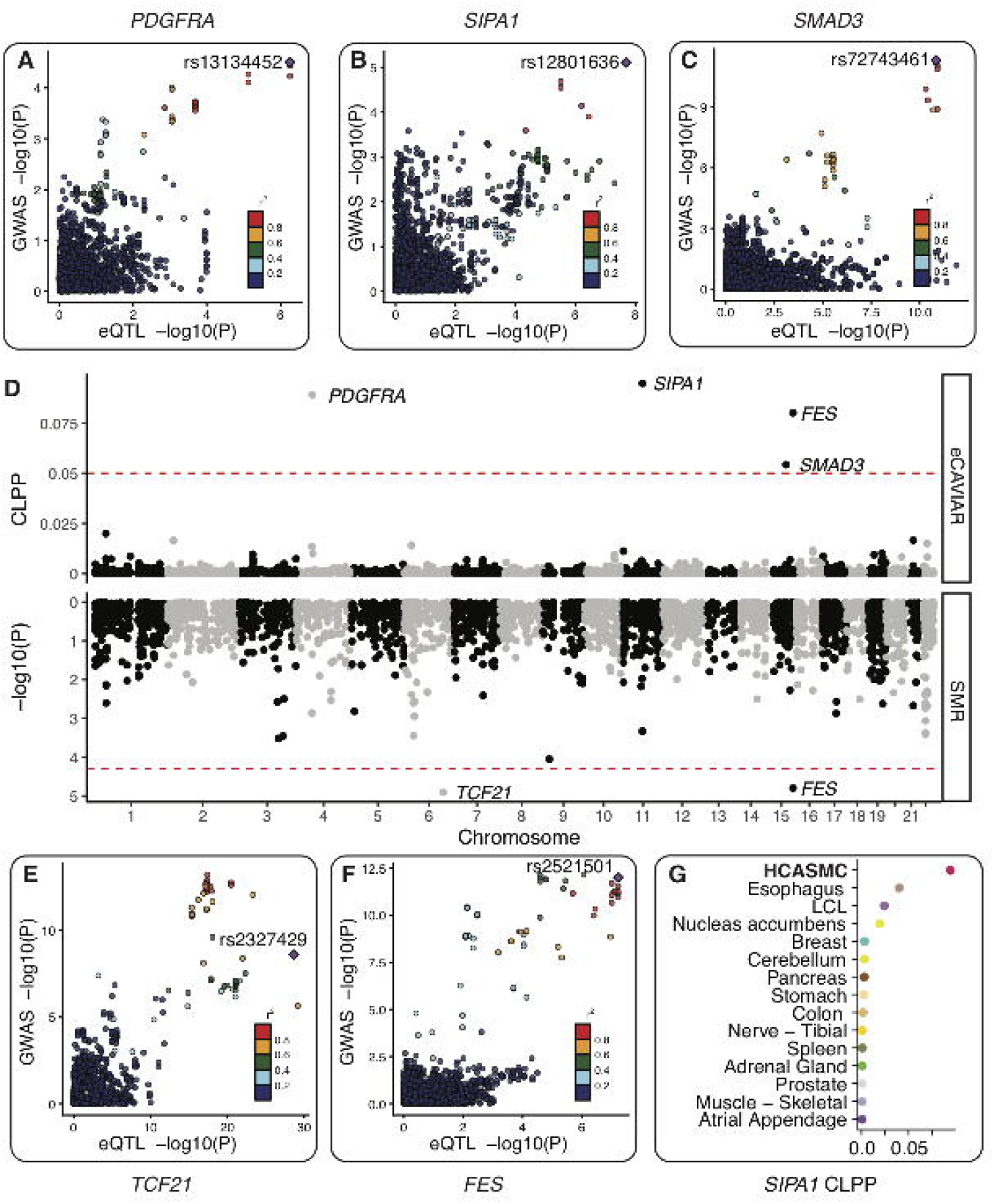
Colocalization between HCASMC eQTL and coronary artery disease GWAS. (**A-C**) Three potential causal genes identified by eCAVIAR. (**A**) Platelet-derived growth factor alpha (*PDFGRA*) eQTL signal colocalized with the *KDR* GWAS locus, which reached FDR < 0.05 significance in the latest CARDIoGRAMplusC4D and UK Biobank GWAS meta-analysis^6^. (**B**) Signal-Induced Proliferation-Associated 1 (*SIPA1*) eQTL signal colocalized with the *PCNX3* locus, which reached genome-wide significance (p-value < 9.71 ×10^−9^) in Howson *et al*.^5^ Note that this study only genotyped selected loci but have a larger sample size than the UK Biobank study. (**C**) *SMAD3* eQTL signal colocalized with the *SMAD3* locus, which is newly identified in the UK Biobank meta-analysis^6^. (**D**) Transcriptome-wide colocalization signals between HCASMC eQTL and CAD GWAS. We used eCAVIAR (top) and SMR (bottom) to fine-map GWAS causal variants and to identify eQTL signals that can explain CAD risk variants (see **Methods**). We found five genes whose eQTL signals show significant colocalization with CAD GWAS signal (SMR FDR < 0.05 or eCAVIAR colocalization posterior probability > 0.05). (**E-F**) Two potential causal genes identified by SMR. (**E**) Transcription factor 21 (*TCF21*) eQTL signal colocalized with the *TCF21* locus, which was identified by Schunkert *et al*.^74^ and replicated in the UK Biobank meta-analysis. (**F**) *FES* eQTL signal colocalized with the *FURIN-FES* locus, which was identified by Deloukas *et al*.^75^ and replicated in the UK Biobank meta-analysis. (**G**) *SIPA1* colocalization is strongest in HCASMC, indicating that this gene influences CAD risk primarily through HCASMC.

**Fig. 4.**
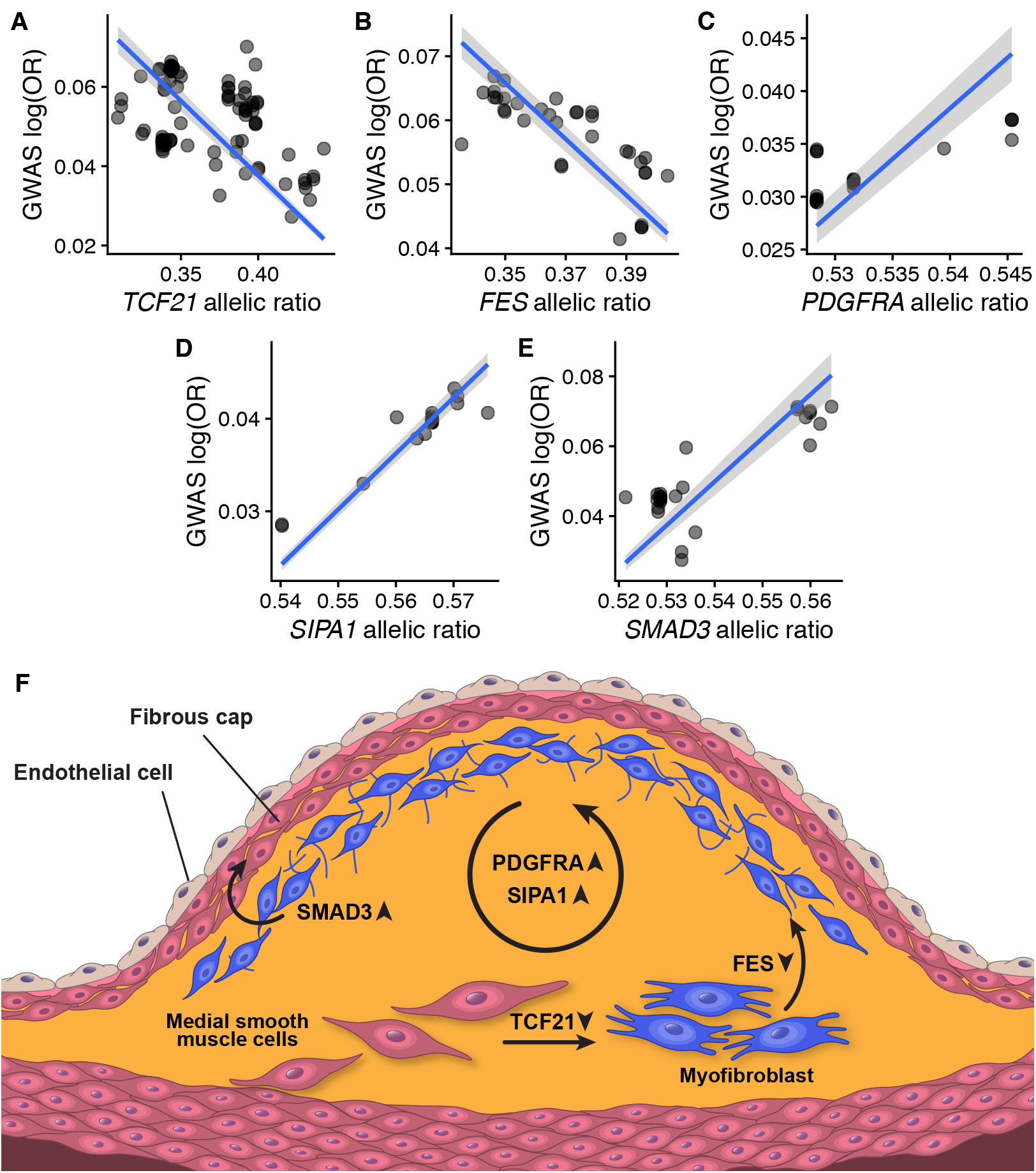
Causal genes are involved in HCASMC-related vascular remodeling. (**A-E**) We determined the direction of effect, i.e. whether gene expression upregulation increases risk, using the correlation between the GWAS and the eQTL study effect sizes on SNPs with p-value < 1×10^−3^ in both datasets. Upregulation of genes *TCF21* and *FES* are protective against CAD risk, and upregulation of *SMAD3, PDGFRA*, and *SIPA1* increases CAD risk. (**F**) Hypothetical functions of five potential causal genes. Upregulation of *TCF21* facilitates the transition of smooth muscle cells from a contractile to a synthetic state^76^. Upon phenotypic transition, *FES* assists in smooth muscle cell migration to the neo-intima^77^. Both *SIPA1* and *PDGFRA* promotes HCASMC proliferation^50,78^. *SMAD3* induces synthetic smooth muscle re-differentiation into the synthetic phenotype for vessel wall repair^79^. Upward arrows indicate genetic upregulation increases CAD risk, and downward arrows indicate genetic upregulation is protective against CAD risk.

## Discussion

In this study, we have integrated genomic, transcriptomic, and epigenetic datasets to create the first map of genetic regulation of gene expression in human coronary artery smooth muscle cells. Comparison with publicly available transcriptomic and epigenomic datasets in GTEx and ENCODE revealed regulatory patterns specific to HCASMC. By comparing against neighboring tissues in GTEx, we found thousands of differentially expressed genes, which were enriched in pathways such as EMT, protein secretion and cellular proliferation, consistent with our current understanding of HCASMC physiology *in vivo*. In comparison with ENCODE, we found 7332 (~2.1%) open chromatin peaks unique to HCASMC, and showed that these peaks are enriched with binding motifs for Forkhead box family proteins, which are known to regulate cell-type-specific gene expression^43^. FOXP1 in particular has been shown to increase collagen production in smooth muscle cells^44^, supporting a potential role in extracellular matrix remodeling in the vessel wall.

Using both transcriptomic and epigenomic profiles, we established that HCASMC represent an important cell type for coronary artery disease. On a tissue-level, we demonstrated that genes highly expressed in HCASMC, coronary artery and adipose tissue are enriched for SNPs associated with CAD risk. While the proximal aortic wall is also susceptible to atherosclerosis, the coronary arteries represent the primary origin of ischemic coronary artery disease in humans^9^. Given that the majority of coronary arteries in the epicardium are encapsulated by perivascular adipose tissue in individuals with disease, one would expect these tissues to share gene responses involved in both vascular inflammation and lipid homeostasis^45^. Further, we demonstrated that HCASMC, endothelial cells, and immune cells also contribute towards the genetic risk of coronary artery disease. Recent -omic profiling of human aortic endothelial cells (HAECs) isolated from various donors identified a number of genetic variants and transcriptional networks mediating responses to oxidized phospholipids and pro-inflammatory stimuli^46^. Likewise, systems approaches investigating resident macrophages and other immune cells involved in vessel inflammation have provided additional insights into context-specific disease mechanisms^47,48^.

Our integrative analyses identified a number of CAD-associated genes that may offer clues into potentially targetable HCASMC-mediated disease mechanisms. Although two of these associated genes, *TCF21* and *SMAD3*, have established roles in regulating vascular remodeling and inflammation during disease^12,16,49^, the other identified genes, *PDGFRA, FES* and *SIPA1*, appear to be novel SMC associated genes. While the role for *PDGFRB* mediated signaling has been well documented in atherosclerosis and modulation of SMC phenotype, the possible involvement of *PDGFRA* has not been investigated in detail^50,51^. Interestingly, *FES* and *SIPA1* were found to harbor CpGs identified in current smokers in the Rotterdam Study, based on targeted methylation profiling of CAD loci in whole blood^52^. The two identified CpGs in *FES* were located near the transcription start site, while the one CpG identified in *SIPA1* was located in the 5’-UTR, suggesting potential environmental influences on gene expression levels. *SIPA1* encodes a mitogen induced GTPase activating protein (GAP), specifically activating Ras and Rap GTPases^53^. *SIPA1* may be a unique mitogen response signal in HCASMC undergoing phenotypic transition in the injured vessel wall; however, these hypotheses should be explored in relevant functional models. Another HCASMC eQTL variant, rs2327429, located in the *TCF21* promoter region, was also the lead SNP in this locus in a recent CAD meta-analysis and has been identified as an mQTL for *TCF21* expression in two separate studies^54,55^. These data suggest that regulation of methylation is a novel molecular trait that may mediate risk for CAD. Splicing QTL colocalization analysis reveals that alternative splicing in *SMG9* also influences CAD risk. *SMG9* has been shown to regulate the non-sense mediated decay (NMD) pathway in human cells, and has been implicated in several developmental disorders such as brain malformations and congenital heart disease^56^.

In summary, the current study confirms the value of detailed genomic and genetic analyses of disease-related tissues and cell types, which when analyzed in the context of publicly available data can provide deep insights into the physiology of human traits and pathophysiology of complex human disease. We expect that these findings will provide a rich resource for the community and prompt detailed functional investigations of candidate loci for preclinical development.

## Supplemental Data

Supplemental Data include fifteen figures and five tables.

## Acknowledgement

We thank Professor Nicolas Mermod for providing biological material and Normal Cyr for making illustrations. B. L. is supported in part by the Stanford Center for Computational, Evolutionary and Human Genetics Fellowship. T.Q is supported by NIH grants R01HL109512(NIH), R01HL134817 (NIH), R33HL120757 (NIH), R01DK107437 (NIH), and R01HL139478 (NIH). C.L.M is supported by R00HL125912 (NIH). S.B.M. is supported by R33HL120757 (NHLBI), U01HG009431 (NHGRI; ENCODE4), R01MH101814 (NIH Common Fund; GTEx Program), R01HG008150 (NHGRI; Non-Coding Variants Program) and the Edward Mallinckrodt Jr. Foundation.

## Web Resources

**Data and code availability**. RNA sequencing data has been deposited at Gene Expression Omnibus (GEO), accession number GSE113348. All eQTL and sQTL summary statistics are accessible through the website http://montgomerylab.stanford.edu/resources.html. All code used to perform analyses and generate figures are in the GitHub repository: https://github.com/boxiangliu/hcasmc_eqtl

**URLs.** GATK, https://software.broadinstitute.org/gatk/; BWA, https://github.com/lh3/bwa; STAR, https://github.com/alexdobin/STAR; Picard, https://broadinstitute.github.io/picard/; RASQUAL, https://github.com/natsuhiko/rasqual; Beagle, https://faculty.washington.edu/browning/beagle/beagle.html; 1000 Genomes, http://www.internationalgenome.org/; VerifyBamID, https://genome.sph.umich.edu/wiki/VerifyBamID; FastQC, https://www.bioinformatics.babraham.ac.uk/projects/fastqc/; WASP, https://github.com/bmvdgeijn/WASP; RNA-SeQC, http://archive.broadinstitute.org/cancer/cga/rna-seqc; GENCODE, https://www.gencodegenes.org/; ENCODE, https://www.encodeproject.org/; Bowtie2, http://bowtie-bio.sourceforge.net/bowtie2/index.shtml; MACS2, https://github.com/taoliu/MACS; ENCODE ATAC-seq/DNAse-seq pipeline, https://github.com/kundaielab/atac_dnase_pipelines; DESeq2, https://bioconductor.org/packages/release/bioc/html/DESeq2.html; sva, https://bioconductor.org/packages/release/bioc/html/sva.html; NOISeq, https://bioconductor.org/packages/release/bioc/html/NOISeq.html; bedtools, http://bedtools.readthedocs.io/en/latest/; JASPAR, http://jaspar.genereg.net/; LD score regression, https://github.com/bulik/ldsc; LeafCutter, https://github.com/davidaknowles/leafcutter; GREGOR, https://genome.sph.umich.edu/wiki/GREGOR; PLINK, https://www.cog-genomics.org/plink2; FastQTL, http://fastqtl.sourceforge.net/; TreeQTL, http://www.bioinformatics.org/treeqtl/; METASOFT, http://genetics.cs.ucla.edu/meta/; SMR, http://cnsgenomics.com/software/smr/#Overview; eCAVIAR, http://zarlab.cs.ucla.edu/tag/ecaviar/; FINEMAP: http://www.christianbenner.com/#.

